# Thyroid hormone induces B cells abnormal differentiation via overexpression of B cell-activating factor

**DOI:** 10.1101/2022.03.07.483380

**Authors:** Shu Liu, Guo-Qing Li, Qing-Wei Gu, Jie Wang, Xin Cao, Yi Zhong, Jing-Jing Miao, Qi Sun, Wen-Sha Gu, Xiao-Ming Mao

## Abstract

Breakdown of tolerance and abnormal activation in B cells is an important mechanism in Graves’ disease (GD) pathogenesis. However, the mechanism by which B cells are abnormal differentiated and activated in GD remains elusive. Here, we show that elevated BAFF expression is positively correlated with serum thyroid hormone (TH) levels in GD patients and high TH levels can induce BAFF overexpression and lead to the abnormal differentiation of B cells in mice. This BAFF overexpression can be seen in many tissues. In the spleens of mice, high TH levels induce M1 macrophages polarization, which generates BAFF overexpression. Our findings open a new perspective on the interactions between endocrine and immune system and provide insight into the involvement of thyroid hormones in the development and progression of GD.

## Introduction

B lymphocytes are essential cells for a functional immune system can also contribute to autoimmune diseases Nearly half of the newly generated B cells are self-reactive, and various selection checkpoints are enforced along B cell development and maturation pathways to increase immune function in hosts while preserving self-integrity (1). The breakdown of B cell tolerance plays a key role in the pathogenesis of many autoimmune diseases (2-5). In Graves’ disease (GD) (6), production of thyroid-stimulating hormone receptor antibody (TRAb) by activated B cell is the main pathogenesis. TRAb persistently stimulates thyroid follicular cells, induces thyroid follicular cell hyperplasia, and secretes excessive thyroid hormones (THs). Unfortunately, the mechanism by which autoreactive B cell escape self-tolerance checkpoints remains largely unknown (7).

B cell-activating factor (BAFF; CD257) belongs to the tumor necrosis factor (TNF)-ligand family, binds to the BAFF receptor (BAFFR) of B cells, and activates several downstream pathways that regulate basic survival functions. These survival functions include protein synthesis and energy metabolism required to extend the half-life of immature, transitional, and mature B cells, leading to increased serum immunoglobulin level (8, 9). BAFF is expressed by monocytes, macrophages, dendritic cells, BM stroma cells, and activated T cells (10), whereas the BAFF receptor is mainly expressed on the surface of B cells. BAFF plays an important role in the immune homeostasis of B cells (11). Excessive BAFF expression is closely involved in the breakdown of B-cell tolerance and autoantibody production, resulting in various autoimmune diseases, including Sjögren’s syndrome (SS), systemic lupus erythematosus (SLE), and rheumatoid arthritis (RA) (12, 13). BAFF overexpression has also been found in GD patients (14, 15) and is associated with thyroid autoantibody production (14).

Although some studies have suggested that BAFF genetic variants are related to abnormal BAFF expression in GD patients, only a small number of GD patients have genetic (16, 17). The reasons for BAFF overexpression in GD patients, as well as whether excessive BAFF expression leads to the breakdown of B cell tolerance, are still poorly understood and need to be elucidated.

The main pathological characteristic of Graves’ hyperthyroidism (GH) is TH overproduction. THs mainly include triiodothyronine (T3) and tetraiodothyronine (T4). TH overproduction affects various organ systems and induces many symptoms, such as palpitations, fatigue, tremor, anxiety, disturbed sleep, weight loss, heat intolerance, sweating, and polydipsia (18). Excess TH levels also influence B-cell activation, differentiation, and proliferation. The high circulating levels of T3 stimulated B cells to form plasma cells (PCs) and increase the proportion of plasma PCs in the bone marrow (BM), which is a long-lived PC (LLPC) survival niche in mice (19, 20). PCs are terminally differentiated B-cells that produce antibodies and provide immediate and long-term protection against pathogens. LLPC overproduction is involved in the development of autoimmune disease (21). However, the mechanisms by which THs induce B-cell proliferation and activation are not fully understood.

In the present study, we found that serum BAFF levels were elevated and positively correlated with FT4 and TRAb, and the proportion of CD86+ and CD80+ B cells was markedly increased in GD patients. To reveal that THs contribute to BAFF overexpression and activate B cells, a mouse model with high-level T3 was used. We showed that high circulating levels of T3 can induce BAFF overexpression and interfere with B cell differentiation. In the spleen of these mice, high T3 levels induce M1 macrophages polarization through interferon gamma (IFN-γ), leading to BAFF overexpression. Our study provides new evidence of endocrine and immune system crosstalk.

## Results

### TH was positively correlated with BAFF expression in GD patients

We recruited 84 patients with GD and 27 healthy donors (general clinical characteristics are summarized in Table 1 and Figure 1A) to evaluate BAFF expression. The results showed that the serum BAFF levels were significantly higher in GD patients than in healthy donors (Figure 1B), which is consistent with results reported in previous studies (14, 15, 22). To determine the factors related to BAFF overexpression, serum FT3, FT4, TRAb, TGAb, and TPOAb levels were measured, and the relationship between these data and BAFF was evaluated. We found that serum BAFF levels were positively correlated with serum FT4 and TRAb levels (Figure 1, C and D). However, there was no significant correlation between BAFF levels and FT3, TGAb, or TPOAb levels (Figure 1, E-G)

**Figure 1.**
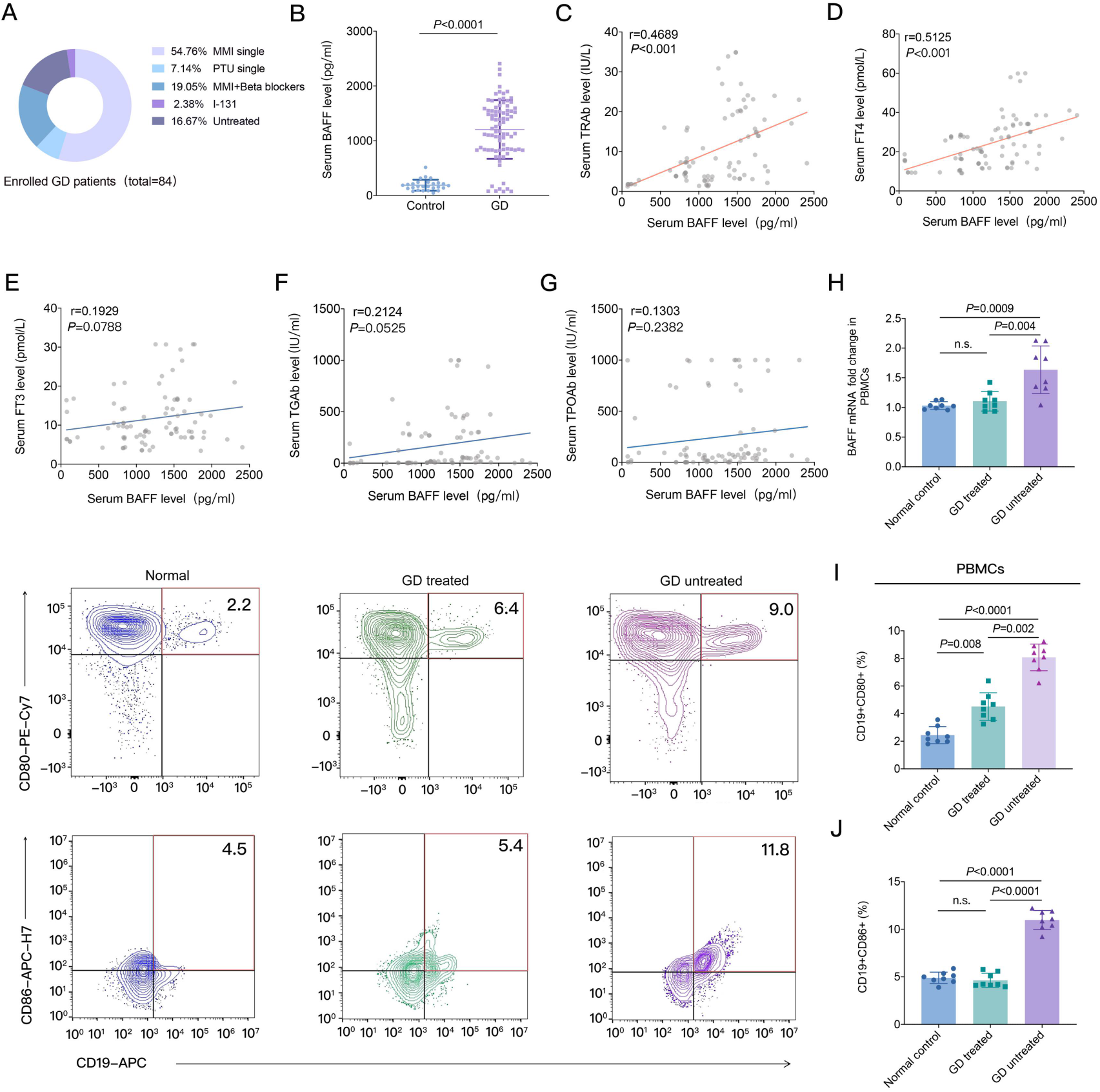
The correlations between serum BAFF and THs, thyroid antibodies, and activation of B cells in GD patients. **(A)** Schematic representation of the enrolled GD patients, in which the proportion of patients treated with antithyroid drugs or I^131^ is 83.33% and untreated is 16.67%. **(B)** Representative of the difference in serum BAFF levels between GD patients (n=84) and healthy donors (n = 27). Data are presented as mean ± S.D. and *P* value between the two groups was calculated by two-tailed unpaired *t*-test. **(C-G)**, The relationship of serum BAFF level with the serum TRAb, FT4, FT3, TPOAb, and TGAb levels in GD patients (n = 84) and healthy donors (n = 27). Correlation coefficient and *P* values are assessed by two-sided Spearman’s rank correlation coefficient test. **(H)** BAFF mRNA expression in the peripheral blood mononuclear cells (PBMCs) of the healthy donors (n = 8), treated (n = 8) and untreated (n =8) GD patients. Data are presented as mean ± S.D. and *P* values are assessed by unpaired *t*-test with Welch’s correction. **(I)** Representative expressions of CD80 and CD86 on CD19+ B cells isolated from the PBMCs of the healthy donors (n=8), treated (n=8) and untreated (n = 8) GD patients. The average fractions of CD19+ B cells positive for CD80 and CD86 are shown at the right panels. Data are presented as mean ± S.D. and *P* values are assessed by two-tailed unpaired *t*-test.

**Table 1.** The general clinical characteristics of GD patients and healthy donors

| Group | Normal<br>Range | Control | Treated GD | Untreated GD | P value |
| --- | --- | --- | --- | --- | --- |
| n | - | 27 | 70 | 14 | - |
| Gender (female/male) (female%) | - | 22/5(81.5%) | 59/11(84.2%) | 12/2 (85.7%) | .895 |
| Age (yr) | - | 49.1±11.7 | 47.0±15.1 | 46.5±12.4 | .960 |
| Smoker (%) | - | 4.0% | 4.8% | 7.1% | .781 |
| Family history of thyroid disease (%) | - | 3.1% | 17.1%<sup>a</sup> | 21.4%<sup>b</sup> | <.001 |
| Other related autoimmune diseases (%) | - | 7.4% | 17.9%<sup>a</sup> | 21.4%<sup>b</sup> | .030 |
| WBC (×10<sup>9</sup>/L) | 3.5-9.5 | 6.09±1.12 | 5.62±1.73 | 5.81±1.48 | .078 |
| ALT (U/L) | 0-40 | 22.10(18.19,28.32) | 36.17(27.50,46.80)<sup>a</sup> | 27.12(17.01,29.37) | <.001 |
| AST(U/L) | 0-40 | 19.06(17.17,23.36) | 29.87(24.76,28.61)<sup>a</sup> | 22.72(18.61,27.95) | <.05 |
| TSH(mIU/L) | 0.35-4.94 | 2.93(2.29,3.80) | 0.023(0.0065,2.71)<sup>a</sup> | 0.0065(0.0065,0.0107)<sup>bc</sup> | <.001 |
| FT3(pmol/L) | 2.63-5.7 | 4.63(3.75,5.12) | 10.61(3.94,31.10)<sup>a</sup> | 21.10(12.65,34.70)<sup>bc</sup> | <.001 |
| FT4(pmol/L) | 9.0-19.0 | 14.22(11.18,17.41) | 22.65(12.50,38.61)<sup>a</sup> | 31.30(19.75,42.08)<sup>bc</sup> | <.001 |
| TRAb(IU/L) | <1.75 | 0.64(0.32,0.89) | 3.77(0.62,8.65)<sup>a</sup> | 10.32(3.15,12.91)<sup>bc</sup> | <.001 |
| TGAb(IU/mL) | <115.0 | 12.49(3.48,27.09) | 80.35(5.86,130.91)<sup>a</sup> | 79.01(6.72,118.87)<sup>b</sup> | .013 |
| TPOAb(IU/L) | <34.0 | 8.52(3.77,17.02) | 276.30(23.95,504.01)<sup>a</sup> | 298.23(18.98,598.23)<sup>b</sup> | <.001 |

Age and WBC presented are mean ± S.D. and statistical significance is assessed by One-way variance analysis. Other clinical data are presented the 25th and 75th percentiles. Statistical significance is determined by the Mann-Whitney or Kruskal-Wallis test. <sup>a,b</sup>, compared with the control. <sup>c</sup>, compared with the treated GD group. Abbreviations: WBC, the actual number of white blood cells; ALT, alanine transaminase; AST, aspartate aminotransferase; FT3, free triiodothyronine; FT4, free thyroxine; TSH, thyroid-stimulating hormone; TR-Ab, thyrotropin receptor antibody; TG-Ab, thyroglobulin antibody; TPO-Ab, thyroid peroxidase.

Both CD80 and CD86 are expressed on activated B cells and are markers of B cell activation (23-25). TRAb production by activated B-cells is the core pathogenesis of GD. To evaluate the effects of THs on BAFF expression and B cell activation, we divided GD patients into two groups: the treated (antithyroid drugs or I^131^ treatment) and untreated groups (Table 1). As expected, TH levels were significantly lower in the treated group than in the untreated group. The proportions of CD80+ and CD86+ B cells and BAFF mRNA expression in peripheral blood mononuclear cells (PBMCs) were significantly increased in the untreated GD group compared with those in the control group (Figure1, H-J). It is interesting to note that following the decline in TH levels in the treated group (Table 1), the BAFF mRNA expression and proportions of CD80+ and CD86+ B cells significantly decreased in the treated group compared with those in the untreated group (Figure, 1H-J). Compared to the control group, the proportion of CD80+ B cells significantly increased in the treated group (Figure, 1I), but there was no significant difference in BAFF mRNA expression and the proportion of CD86+ B cells in the treated group (Figure 1H and I).

### TH induced BAFF overexpression in T3 treated mice

To evaluate the effects of THs on BAFF expression, we used 5 μg/10 g of T3) to create a high-level T3 mouse model, BAFF shRNA (Bs) to interfere with BAFF expression, and clodronate liposomes (CLs) to eliminate macrophages in the mice (Figure 2A). After six weeks of continuous subcutaneous injection of T3, the average body weight decreased, while the average food and water consumption increased in all three T3 groups compared with that in the control groups (Figure 2, B-D). The serum T3 levels markedly increased in the T3, T3+Bs, and T3+ negative control shRNA (NCs) groups of mice compared with those in the control group, exhibiting more than a two-fold increase (Figure 2E). Meanwhile, the serum BAFF levels and BAFF mRNA expression in bone marrow (BM) and spleen significantly increased in the T3 and T3+NCs groups compared with those in the control and T3+Bs groups, respectively (Figure 2,F-H). Furthermore, BAFF protein expression in the spleen also significantly increased in the T3 and T3+NCs groups compared with that in the control and T3+Bs groups (Figure 2I).

**Figure 2.**
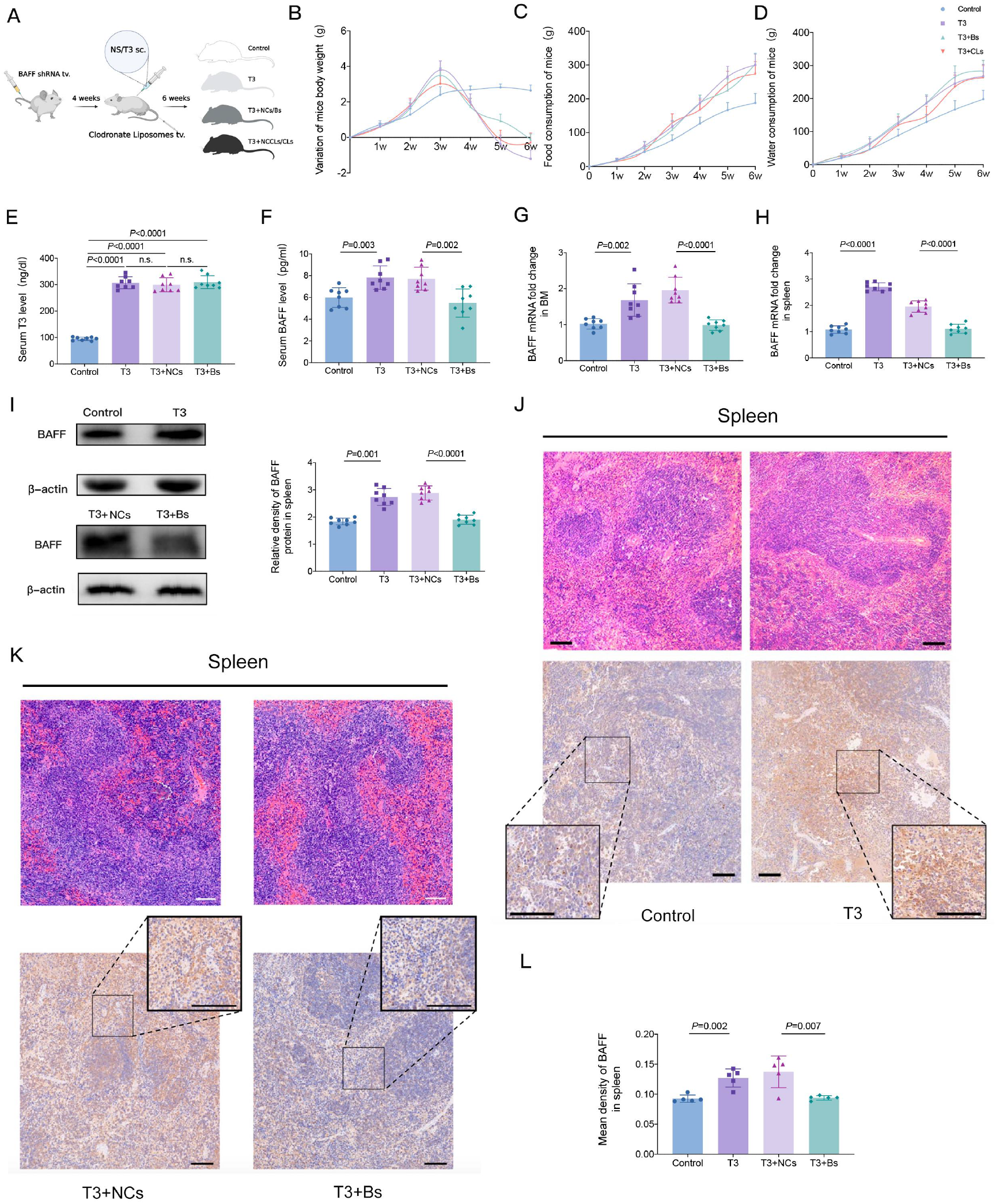
BAFF overexpression induced by TH and the results of inhibiting BAFF expression by shRNA in mice. **(A)** Schematic representation of the experimental protocol. The mice were divided six groups: Control, T3, T3+negative control shRNA (T3+NCs), T3+BAFF shRNA (T3+Bs), T3+negative clodronate liposome control (PBS) (T3+NCCls), and T3+clodronate liposome (T3+CLs) groups. The mouse of T3 group was treated with 5 μg/10 g T3 for 6 weeks. The mice of T3+BS and T3+ NCs groups were treated with shRNA and negative control shRNA for 4 weeks before 6 weeks of T3 treatment. **(B-D)** Representative the changes of body weight **(B)**, food consumption **(C),** and water consumption **(D)** in mice of Control, T3, T3+NCs and T3+Bs groups during the 6 weeks. **(E-H)**, Representative the changes of serum T3 level **(E)**, serum BAFF level **(F)**, BAFF mRNA expression in BM **(G)** and BAFF mRNA expression in spleen **(H)** for the four groups. (n = 8 biological replicates for each group of the mice). Data are presented as mean ± S.D. Statistical significance is assessed by two-tailed unpaired *t*-test. **(I)** Representative the changes of BAFF protein expression in mice spleen of the four groups. Data are presented as mean ± S.D. Statistical significance is assessed by two-tailed unpaired *t*-test. **(J** and **K)** The spleen pathological sections stained with hematoxylin and eosin (top panels), and immunohistochemically stained with anti-BAFF antibody (lower panels) for the four groups. **(L)** Representative the fraction of BAFF positive cells in the four groups. The cytoplasm of BAFF-positive cells was visualised as a brownish yellow stain using a pathological image analysis system. Five visual fields were randomly selected from each section. Integral absorbance was calculated as positive BAFF expression (n = 5 biological replicates). Mean density was calculated by mean IOD/ mean area. Scale bar = 100 um. Data are presented as mean ± S.D. Statistical significance is assessed by two-tailed unpaired *t*-test.

We noted that the spleen white pulp compartment was expanded and fused in T3-treated mice, as analyzed by H&E staining (Figure 2J). Immunohistochemical staining confirmed that BAFF expression increased in the spleen of T3-treated mice compared with that in control mice (Figure 2J and L). When shRNA was used to inhibit BAFF expression, the spleen white pulp compartment and fused degree in white pulp areas were reduced in T3+Bs group by H&E staining compared to the T3+NCs group (Figure 2K). Immunohistochemical staining showed that BAFF expression was also reduced in the T3+Bs group compared with the T3+NCs group (Figure 2, K and L).

### Elevated BAFF levels induce abnormal B-cell differentiation

B cell development in human BM proceeds through successive stages to form transitional B cells with low avidity to self-antigens, Immature B cells are formed with the cell surface expression of IgM (26, 27), leave the BM, enter circulation, and migrate to the spleen, completing the early steps of B cell development (28), and then form mature B cells that express IgM and IgD. BAFF is an essential survival factor for splenic B cells and is required for normal splenic B cell numbers (29). Excess of BAFF leads to severe B-cell hyperplasia (30) and promotes the development of autoreactive B-cells and autoantibody production (31, 32).

We assessed whether BAFF overexpression induced by T3 affects B cell differentiation in the BM and spleen. The flow cytometry results showed that the proportion of B220+IgM+ B cells and plasma cells (PCs) in the BM (Figure 3, A-C).) and B220+IgD+IgM+ B cells and PCs in the spleen (Figure 3, A, D and E) were markedly increased in the mice of T3 treated groups compared with those in the mice of the control group. When shRNA was used to interfere with BAFF expression, the proportions of spleen B220+IgM+ B cells and PCs in the BM (Figure 3, A-C), and B220+IgM+IgD+ B cells and PCs in the spleen (Figure 3, A, D and E) were all significantly reduced in the mice of the T3+Bs group compared with those of the T3+NCs group.

**Figure 3.**
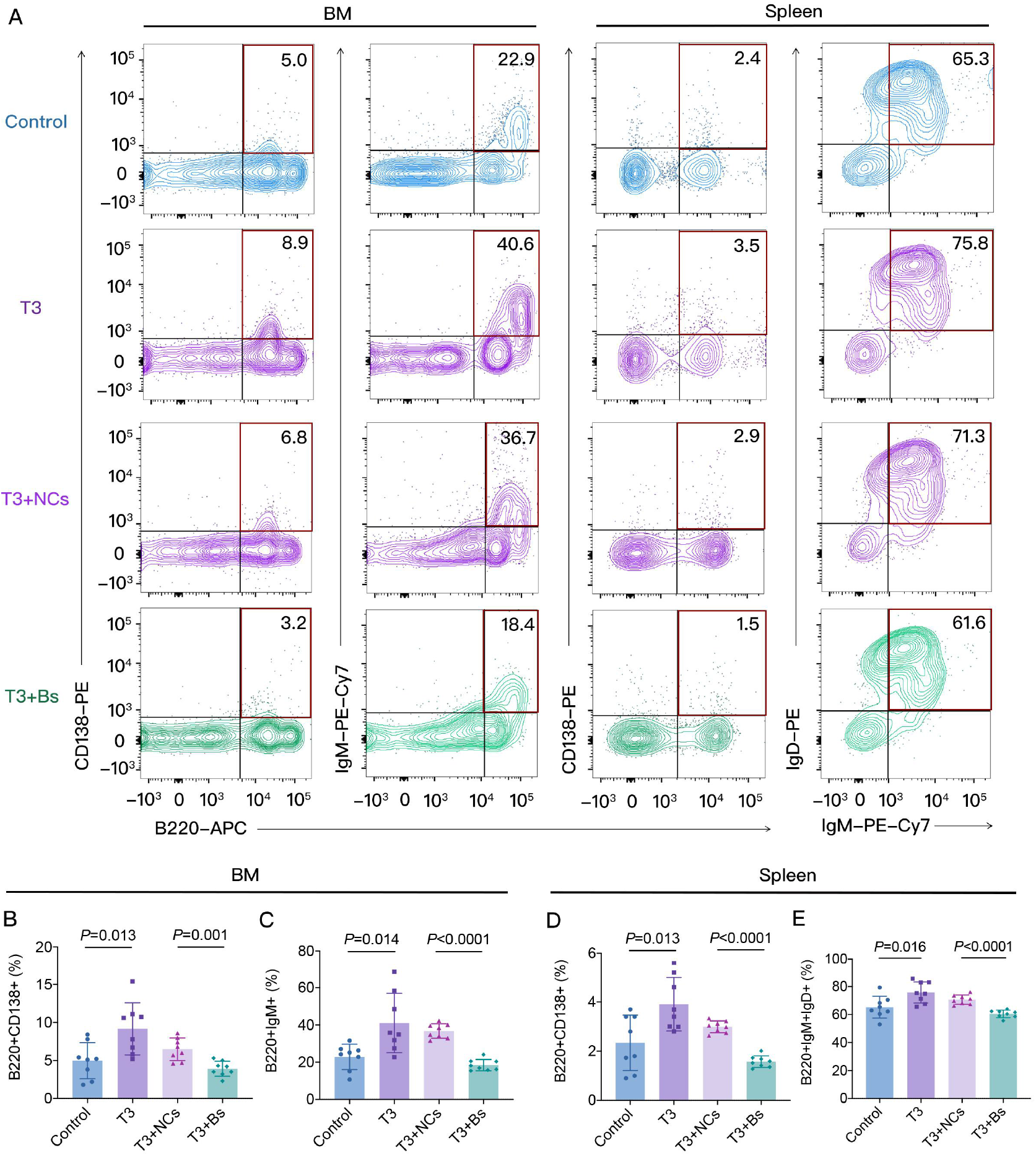
B cells differentiation in BM and spleen of the mice after T3 treatment. **(A)** Representative expressions of CD138 and IgM on B220+ B cells in BM, CD138, IgM and IgD on B220+ B cells in spleen of the control, T3, T3+NCs, and T3+Bs groups**. b-d**, Representative the average fraction of B220+ B cells positive for the CD138 and IgM in BM, and CD138, IgM and IgD in spleen of the four groups. Data are presented as mean ± S.D. (n = 8 independent biological experiments). Statistical significance is assessed by two-sided independent *t-*test.

### High TH levels induce macrophages to express BAFF and polarize them to M1 cells

We found that high T3 levels could induce BAFF overexpression in the BM and spleen of the mice. To further clarify the sources of the overexpression BAFF in the peripheral blood of both humans and mice induced by T3, the thyroid, spleen, muscle, liver, small intestine, lung, brain, kidney, stomach, heart, and pancreas were isolated in the mice after 6 weeks of T3 treatment. BAFF protein expression was significantly increased in most tissues, except for the muscle and stomach, compared with that in the control group (Figure 4A).

**Figure 4.**
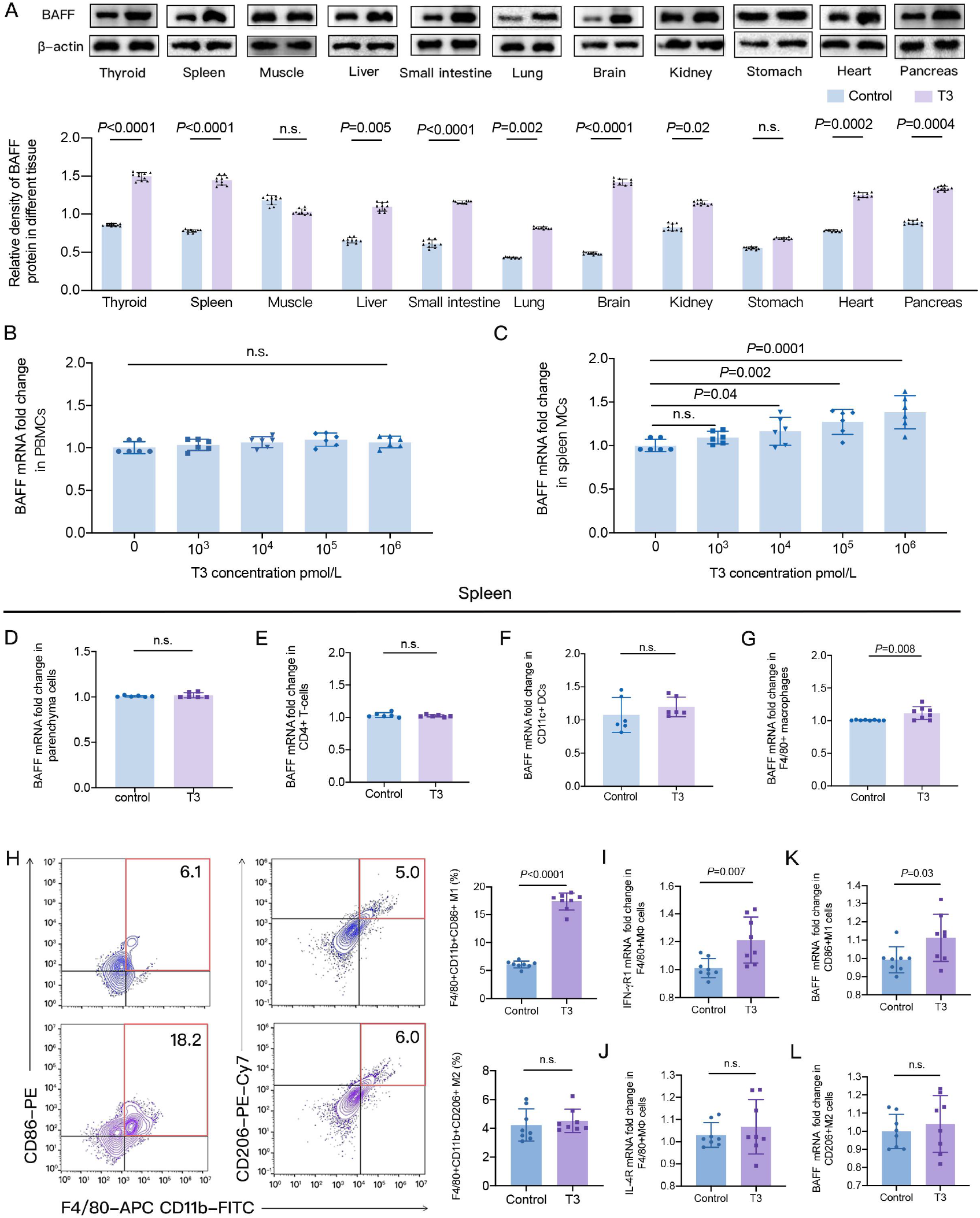
BAFF expression in some tissues and spleen macrophages after T3 stimulation. **(A)** Representative BAFF protein expressions in various tissues of the mice treated with T3 for 6 weeks. The average BAFF protein expressions in various tissues are shown at the bottom panel. Data are presented as mean ± S.D. and *p* value between T3 and control groups is calculated by two-sided independent *t*-test. (n = 8 independent biological experiments). **(B)** Representative BAFF mRNA expression in PBMCs of the normal mice after various concentrations of T3 stimulation. Data are presented as mean ± S.D. and *P* value between 0 and various concentrations of T3 was calculated by two-sided independent *t*-test. (n = 8 independent biological experiments). **(C)** Representative BAFF mRNA expression in spleen MCs of the normal mice after various concentrations of T3 stimulation. Data are presented as mean ± S.D. and *P* value between 0 and various concentrations of T3 is calculated by two-sided independent *t*-test. (n = 8 independent biological experiments). **(D-G)** Representative BAFF mRNA expression of spleen parenchyma cells **(D)**, CD4+T cells **(E)**, CD11c+ DC cells **(F)** and macrophages **(G)** in the mice treated with T3 for 6 weeks and control. Data are presented as mean ± S.D. and *P* value between control and T3 groups is calculated by two-sided independent *t*-test. (n = 8 independent biological experiments). **(H)** Representative expressions of CD86 on spleen F4/80+ CD11b+ macrophages in the mice spleen after 6 weeks T3 treatment. The average fraction of spleen F4/80 CD11b macrophages positive for CD86 is shown at the right panel. Data are presented as mean ± S.D. and *P* value is assessed by two-sided independent *t*-test. (n = 8 independent biological experiments). **(I)** Representative expressions of CD206 on spleen F4/80+CD11b+ macrophages. The average fraction of spleen F4/80 CD11b macrophages positive for CD206 is shown at the right panel. Data are presented as mean ± S.D. and *P* value is assessed by two-sided independent *t*-test. (n = 8 independent biological). **(J** and **K)** Representative mRNA expression of IFN-γR1 **(J)** and IL-4 **(K)** in MФ macrophages, Data are presented as mean ± S.D. *P* value is assessed by two-sided independent *t*-test. (n = 8 independent biological experiments). **(M** and **N)** Representative mRNA expression BAFF mRNA expression in M1 macrophages **(M)** and M2 macrophages **(N)**. Data are presented as mean ± S.D. *P* value is assessed by two-sided independent *t*-test. (n = 8 independent biological experiments).

Although BAFF can be expressed by different immune cells, including monocytes, macrophages, dendritic cells, neutrophils, follicular dendritic cells, epithelial cells, and stromal cells (33-37), it is not known which cells can be induced by T3 to overexpress BAFF. There is a distinction in the components between peripheral blood and tissue mononuclear cells, the macrophages rarely are found in peripheral blood. We used various concentrations of T3 to stimulate peripheral blood mononuclear cells (PBMCs) and spleen mononuclear cells (MCs) for 72 hours, it is interesting that followed by the concentrations of T3 elevation (10^4^, 10^5^, 10^6^ pmol/L), the BAFF mRNA expression significantly increased in spleen MCs, but T3 did not affect BAFF expression in PBMCs (Figure 4, B and C). To confirm the reliability of our results, we used T3 at various concentrations for 72 h to stimulate PBMCs. No apparent difference in PBMC apoptosis was observed (Supplemental Figure 1).

To clarify which cells can be induced by T3 to overexpress BAFF in the spleen, we used 5 μg/10 g of T3 to treat the mice for six weeks and isolated spleen parenchyma, DCs, T cells and macrophages. The results showed that only macrophages highly expressed BAFF mRNA after T3 stimulation (Figure 4, D-G). The heterogeneity of macrophage phenotypes is commonly referred to as polarization that conventionally subdivides macrophages into three groups: naïve (Mφ; also called M0), which readily differentiate into pro-inflammatory (M1) and anti-inflammatory (M2) (38, 39) phenotypes. M1 macrophages are activated by toll-like receptor ligands, such as lipopolysaccharide (LPS) and interferon gamma (IFN-γ), which express proinflammatory cytokines, mediate defense of the host from a variety of bacteria, protozoa, and viruses, and have roles in antitumor immunity. M2 macrophages are stimulated by interleukin (IL)-4 or IL-13, have anti-inflammatory and pro-tumoral functions and regulate wound healing (40,41).

We used flow cytometry and RT-PCR to identify the polarization of macrophages and the macrophage subsets in the spleen that can be induced to overexpress BAFF by high T3 levels. The results showed that the proportion of spleen F4/80+CD11b+CD86+ cells (M1) significantly increased, but there was no significant difference in F4/80+CD11b+CD206+ cells (M2) in T3 treated mice compared with those in the control (Figure 4H). The IFN-γR1 and BAFF mRNA expression in spleen F4/80+ cells (Mφ) and M1 cells significantly increased, but IL-4 and BAFF mRNA expression was not significantly different in F4/80+ cells (Mφ) and M2 cells in T3 treated mice, compared with the control (Figure 4, J-M).

### The elimination of macrophages reduces BAFF expression induced by T3

To confirm the effect of T3 on BAFF expression in spleen macrophages, we used clodronate liposome (CLs) to eliminate macrophages in T3 treated mice. The results showed that the spleen weight was similar between the T3 and T3+CLs groups (Figure 5A). Compared with the T3+(negative clodronate liposome control (NCCLs) group, the serum T3 level was not significantly different in the T3+CLs group, but the serum BAFF, spleen BAFF mRNA, and protein expression were all significantly decreased in the T3+CLs group, compared with the T3+NCCLs group (Figure 5, B-E). Furthermore, the spleen IFN-γR1 protein expression was significantly increased, but there was no significant difference in IL-4R protein expression in the T3+CLs group compared with the T3+NCCLs group ((Figure 5F). The immunohistochemical staining results showed that the spleen BAFF protein expression significantly decreased in the T3+CLs group compared with T3+NCCLs group (Figure 5G). The immunofluorescence staining results suggested that both M1 and M2 cells had significant co-localization with BAFF in the NCCLs group. After eliminating macrophages with CLs, only M1 cells colocalized with BAFF (Figure 5, H and I).

**Figure 5.**
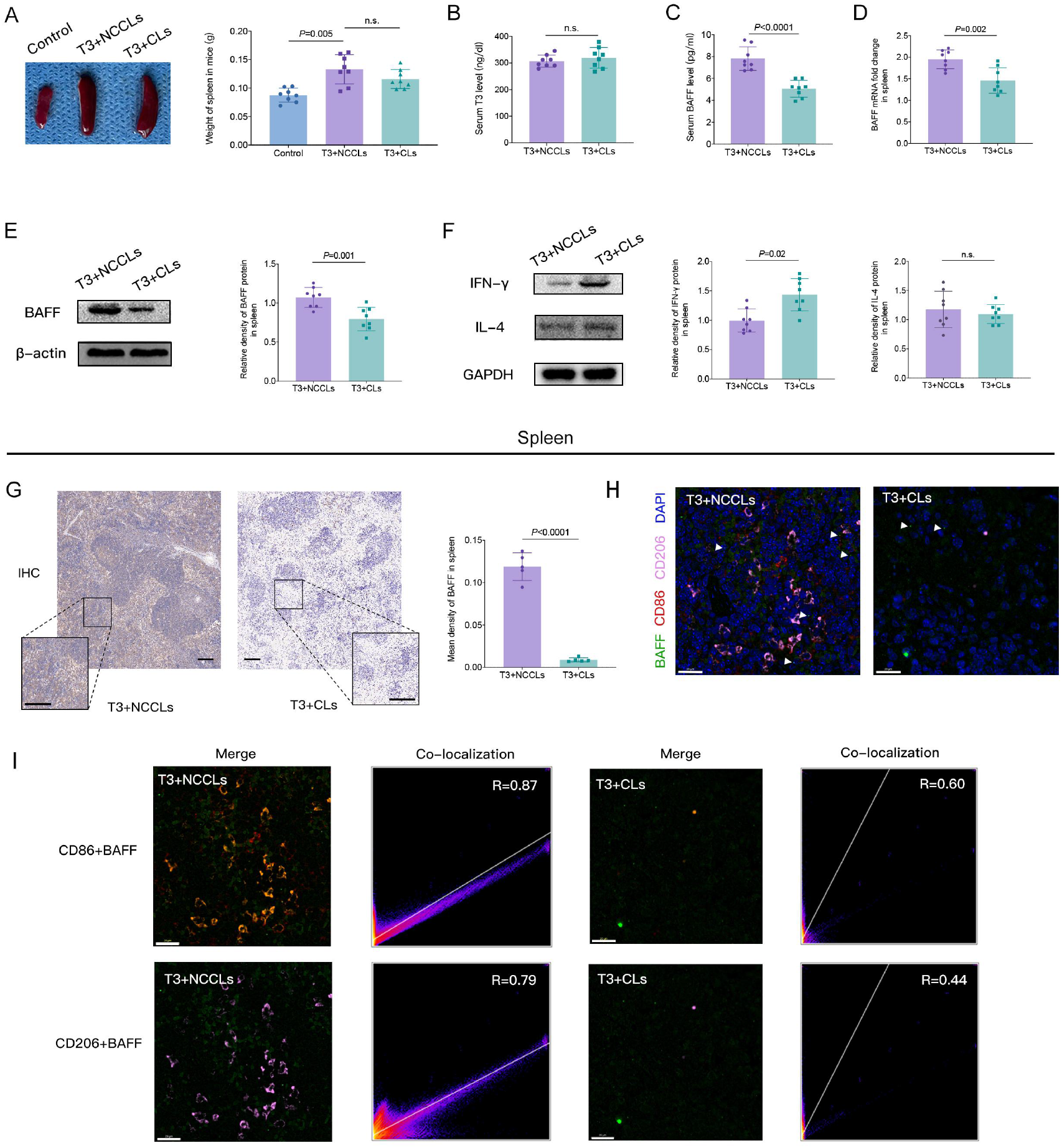
Polarization of spleen MФ macrophages to M1 macrophages by TH stimulation and effects of eliminating macrophages BAFF expression. **(A)** Representative spleen weight changes among the control, T3, T3+NCCls, and T3+Cls groups of mice. The T3+NCCLs and T3+CLs groups were intravenously injected PBS or CLs via tail veins, respectively. Two days later, the mice were injected subcutaneously with T3 (5 μg/10 g) every day for 6 weeks, meanwhile the mice were intravenously injected PBS or CLs via tail veins once a week. The average spleen weights of the four groups of mice are shown at the right panel. Data are presented as mean ± S.D. and *P* value is assessed by two-sided independent *t*-test. (n = 8 independent biological experiments). **(B-D)** Representative the deference of serum T3 (B) and serum BAFF levels **(C)**, and BAFF mRNA expression **(D)** in PBMC between the T3+NCCls and T3+CLs groups. Data are presented as mean ± S.D. and *P* value is assessed by two-sided independent *t*-test. (n = 8 independent biological experiments). **(E** and **F)** Representative the deference of IFN-γR1 **(E)** and IL-4 **(F)** protein expression between the T3+NCCls and T3+CLs groups. Data are presented as mean ± S.D. and *P* value is assessed by two-sided independent *t*-test. (n = 8 independent biological experiments). **(G)** Representative the fraction of BAFF positive cells in T3+NCCls and T3+CLs groups. The cytoplasm of BAFF-positive cells was visualised as a brownish yellow stain using a pathological image analysis system. Five visual fields were randomly selected from each section. Integral absorbance was calculated as positive BAFF expression (n=5 biological replicates). Mean density was calculated by mean IOD/ mean area. Scale bar = 100 um. Data presented are mean ± S.D. Statistical significance is assessed by two-tailed unpaired *t*-test. **(H)** Comparison of co-immunostaining for anti-BAFF (green; white arrows), anti-CD86 (red), anti-CD206 (pink), and DAPI (blue) in spleen between the T3+CLs and T3+NCCLs groups. **(I)** Co-localization of merged images and scatter plots of Pearson coefficient in spleen with anti-BAFF (green), anti-CD86 (red) and anti-CD206 (pink), respectively. The ordinate is the green signal channel, and the abscissa is the red signal channel and the pink signal channel. Five visual fields were randomly selected from each section. Scale bar = 20um. Statistical significance is assessed by Coloc 2.

## Discussion

BAFF promotes the activation and survival of B cells and has been linked to pathogenic B cell responses that underlie autoimmunity and lymphoid hyperplasia in humans and mice (42). Immature B cells in the BM are prone to apoptotic selection at two differentiation stages, associated with IgH chain gene rearrangement and B cell receptor expression, respectively, which indicates that selection by apoptosis is a primary mechanism for eliminating aberrant or autoreactive B cells (43). BAFF can suppress apoptosis of IgM+ B cells, and excess BAFF promotes the survival and maturation of low-affinity self-reactive transitional B cells (8, 44, 45). Transgenic mice overexpressing BAFF develop autoimmune disorders characterized by B cell hyperplasia and autoantibody production including anti-DNA and rheumatoid factor, and eventually succumbed to an immune complex-mediated, lupus-like nephritis (46). B cells were primarily identified for their key role as enhancers of the immune response in autoimmunity because they give rise to autoantibody-producing PCs (47). In addition, blockade of BAFF with the BAFF-specific receptor-Fc can lead to significant reductions in hyperthyroidism in a GD murine model (48). However, apart from some gene variants associated with elevated BAFF expression, little is known about the mechanism of BAFF overexpression in autoimmune disorders, including GD (14-17).

In this study, we discovered that serum BAFF levels are elevated in patients with GD, especially in untreated GD patients treated with antithyroid drugs (ATDs), and serum BAFF levels positively correlate with serum T4, and B cells are activated in our clinical study. High T3 levels can induce BAFF overexpression and promote B cell differentiation into PCs in mice. PCs are terminally differentiated B-cells that produce antibodies that provide immediate and long-term protection against pathogens. Circulating high-affinity antibodies have been sustained for decades by long-lived plasma cells (LLPCs) that mainly reside in the BM (49). BAFFs provide a survival niche for PCs that live in the BM and spleen (50, 51). Previous studies have suggested that LLPCs are an essential component of immunological memory that contributes to the chronicity of antibody-mediated autoimmune diseases(21). In the present study, we found that high T3 levels decreased the proportion of B220+IgG+ cells and PCs in the peripheral blood (Supplemental Figure 2), although peripheral blood BAFF expression was higher in T3 treated mice. This result was consistent with a previous study in which the serum IgG levels did not increase despite the elevated proportion of PCs in the BM and spleen after T3 treatment in mice (19). This might be due to B cells homing to the BM, spleen (52), and tissue (53).

GD is frequently associated with other autoimmune diseases such as vitiligo, rheumatoid arthritis, multiple sclerosis, Sjögren’s disease, systemic lupus erythematosus (54, 55), etc. However, the mechanisms underlying these associations remain unknown. Activation of BAFF plays an important role in the development of these autoimmune diseases (17). In the present study, we found that high TH levels can induce BAFF overexpression, which may explain these associations in patients with GH. Furthermore, ATDs have long been used to treat hyperthyroidism, and the main role of these drugs is to inhibit thyroid peroxidase, thus blocking TH synthesis. The aim of ATD treatment is to reduce TH levels to the normal range in patients with hyperthyroidism. After 12–18 months of ATD treatment in GD patients, more than half of the patients may experience a relapse of hyperthyroidism (56). Our study supports the idea that long-term control of THs with ATDs can reduce GD relapse of GD (57, 58), which may account for the decline in TH and BAFF levels.

Although we did not find a positive correlation between serum FT3 and BAFF in our clinical research, we still selected T3 to study TH with BAFF overexpression and B cell differentiation in mice. T4 is a pro-hormone that exclusively synthesizes and secretes thyroid follicular cells and enters circulation. T4 in circulation reaches target tissues where it is converted into T3 by type 2 iodothyronine deiodinase (DIO2) and plays physiological roles in the tissues. Thus, T3 in circulation only partially reflects this. Furthermore, at the target cell level, the genomic action of THs requires T3 to bind to its specific nuclear receptors (TRs) (59). The lack of a positive correlation between serum FT3 and BAFF may be due to the majority of T3 existing tissues and the circulating free FT4 concentration being fourfold higher than FT3. The relatively small number of volunteers in this study may account for this result.

We found that T3 can induce a large amount of tissue to express BAFF, and only macrophages can be stimulated by T3 to overexpress BAFF in mouse spleens. THs can promote IFN-γR1 upregulation in macrophages and increase IFN-γ expression from Th1 cells (60) that induce MФ polarization to M1, like to T3 induced Bone marrow MФ polarization of bone marrow to M1 (61). The classic genomic actions of THs are mediated by thyroid hormone receptor TRs, and there are two types of TRs encoded by the TRα and TRβ genes. Macrophages express mRNAs of the two major TR isoforms, TRα1 and TRβ1, but only TRβ1 was detected at the protein level^61^. The intracellular action of THs is regulated by the amount of local T3 available for receptor binding (62, 63). The exact mechanism by which THs induce macrophages to express BAFF requires further investigation.

Taken together, our results suggest that THs can induce BAFF overexpression in macrophages to activate B cells and promote B cell differentiation into PCs, which may be involved in the development and progression of GD. Our study provides new evidence of cross-talk between the endocrine and immune systems.

## Methods

### Patients

A total of 84 patients with GD and 27 healthy donors were recruited at Nanjing First Hospital between 2020 and 2021. All patients satisfied the diagnostic criteria for GD, including clinically and biochemically verified hyperthyroidism and a positive thyrotropin receptor antibody (64, 65). The clinical evaluation included patient history, physical examination, and thyroid ultrasonography. Laboratory and diagnostic testing included the determination of serum free thyroxine (FT4) and T3 (FT3) and sensitive TSH (s-TSH) levels, and serum hyrotropin receptor antibody levels. Patients with a low number of leukocytes (less than 3.5×10^9^), subacute thyroiditis, hyperfunctioning thyroid nodules, iodine hyperthyroidism, drug-induced hyperthyroidism, and other causes of hyperthyroidism were excluded. All patients and healthy donors were excluded because of autoimmune, infectious, or other systemic diseases and cancer.

### Mice

C57BL/6J mice at 6–8 weeks of age with an initial weight of 16–22 g were purchased from Beijing Weitong Lihua Experimental Animal Technology Co., Ltd., and maintained under specific pathogen-free conditions (Experimental Animal Center of Nanjing First Hospital, Nanjing Medical University). Animals were bred and housed in individual cages with free access to standard laboratory water and chow.

### Experimental design and specimen preparation

The mice were randomly assigned to control, T3, T3+negative control shRNA (T3+NCs), T3+BAFF shRNA (T3+Bs), T3+negative control liposomes (PBS) (T3+NCCLs), or T3+clodronate liposomes (T3+CLs) groups. The tail veins of the Bs+T3 and T3+ NC groups were intravenously injected with BAFF shRNA or negative shRNA, respectively (GeneChem Co., Ltd., Shanghai, China). Four weeks later, the T3+ NC +T3 and B +T3 groups were injected subcutaneously with T3 (5 μg/10 g) (Meilun Biotech Co., Ltd., Dalian, China) every day for six weeks, and the control group was subcutaneously injected with the same volume of saline. The T3+ NCCL and T3+ CL groups were intravenously injected with PBS or CLs (Liposoma B.V., Amsterdam, Netherlands) via the tail veins. Two days later, the mice were injected subcutaneously with T3 (5 μg/10 g) every day for six weeks, while the mice were intravenously injected with PBS or CLs via the tail veins once a week. This procedure ensures that macrophage subsets are completely eliminated (66, 67).

The mice were tagged with toe clipping and weighed daily before and after administration. The daily amounts of feed and water were recorded. One day post experimental methods, the mice were sacrificed under chloral hydrate anaesthesia, and the blood, bone marrow, spleen, and other tissues were collected for further analyses, as described below.

### Lentivirus production and in vivo infection

Lentivirus construction encompassing BAFF was generated using the AdMax (Microbix) and pSilencer™ adeno 1.0-CMV (Ambion) systems, according to the manufacturer’s recommendations (Genechem Co., Ltd, Shanghai, China). The viruses were packaged, amplified in 293T cells, and purified. Titering was performed on 293T cells using the Adeno-X Rapid Titer kit, according to the manufacturer’s instructions. The anti-BAFF shRNA sequence was CGGGAGAATGCACAGATTT (Shanghai Genechem Co. Ltd., Shanghai, China).

For *in vivo* infection, lentiviruses were injected via the tail vein in 100 µL of PBS containing 7.6×10^7^ IFUs of loaded lentivirus per mouse as described previously (68, 69). The shRNA dose was 10 mg/kg per mouse. A single injection of shRNA was performed, and the mice received daily subcutaneous administration of T3 as described previously for six weeks.

### Sample collection, cell isolation, and histological staining

Peripheral blood samples were collected from GD patients and healthy donors. Serum TSH, FT3, FT4, TGAb, TPOAb, and TRAb levels were measured using a chemiluminescence assay (Centaur XP automated chemiluminescence immunoassay analyzer, Siemens, Germany). Human peripheral blood mononuclear cells (PBMCs) were prepared from fresh peripheral whole blood using Ficoll-Hypaque density gradient centrifugation (TBD Science). CD19+ B cells were isolated from PBMCs by using CD19+ B cell isolation kits (Miltenyi Biotec).

BM cells from mice were obtained by flushing the femurs of animals with culture medium injected through a 21-gauge needle. The thyroid, spleen, muscle, liver, small intestine, pancreas, lung, brain, kidney, stomach, and heart tissues were removed before being smashed on a nylon membrane in a Petri dish with D-hanks. The cell suspensions were centrifuged for 5 min at 200 × *g*, and the cell pellet was resuspended in 3 ml of saline or culture medium supplemented with 10% FBS. The methods used for isolating PBMCs and B cells in mice were the same as those used in humans. CD11c+ DCs, F4/80 macrophages, and CD4+ T cells were isolated from mouse spleen PBMCs using CD11c+ DC cell isolation kits, F4/80 macrophage isolation kits, and CD4+ T cell isolation kits (Miltenyi Biotec).

For histological analysis, the spleen and thyroid were sliced into small pieces and maintained in 4% formalin overnight. The material was dehydrated in graded ethanol baths for 15 min and washed in xylene thrice for 15 min before paraffin embedding. Paraffin blocks with spleens were sliced and 5 mm tissue sections were stained with Harris’s hematoxylin and eosin (H&E staining).

### Immunohistochemical staining of spleen and thyroid tissue

Spleen and thyroid samples were preserved in 10% formalin, dehydrated and embedded in paraffin following routine methods. Paraffin sections were routinely dewaxed, incubated at room temperature with 3% H_2_O_2_, repaired with 0.01 mol/L citric acid solution at high temperature; anti-BAFF antibody (Abcam plc., UK) was used for immunohistochemical staining, and the DAB-H2O2 colorant was added for color rendering. Finally, hematoxylin was used for dehydration and sealed with neutral gum for observation under a light microscope. The cytoplasm of BAFF-positive cells was brownish yellow. A pathological image analysis system was used in the present study. The antibody signal was quantified using the Imageproplus 6.0.

### T3 treatment in vitro

PBMCs and MCs from spleen were isolated and treated with T3 10^3^, 10^4^, 10^5^, 10^6^, and 10^7^ pmol/L (Dalian Meilun Biotechnology Co., Ltd.) for 72 h in a 5% CO2 incubator at 37 °C. Apoptosis of PBMCs was determined by staining with Annexin V-PE (KGA1017, Keygen Biotech).

### Measurement of serum T3 and BAFF levels

Blood was collected from the mouse heart and centrifuged at 250 g for 5 min. Serum was stored at −80 °C until use. The concentration of T_3_ in serum was determined by radioimmunoassay, and the concentration of BAFF in serum was determined by ELISA (Research & Diagnostics Systems, Inc., USA), which was operated strictly according to the manufacturer’s instructions.

### Western blotting

Total tissues were homogenized and total proteins were extracted using a tissue and cell total protein extraction kit (Jiangsu KeyGEN BioTECH Corp., Ltd, Nanjing, China), which was used to analyze the expression of BAFF. Equal amounts of protein were separated by 15% SDS-PAGE and then transferred to polyvinylidene fluoride (PVDF) membranes. Subsequently, the membranes were blocked with 5% nonfat milk for 2 h, then incubated with anti-BAFF (Abcam plc., UK), anti-β-actin (Cell Signaling Technology Inc., Danvers, MA, USA), anti-IFN-γ (Abcam plc., UK), anti-IL-4 (Proteintech Group, Inc., Rosemont, IL, USA), and anti-GAPDH (Abcam plc., UK), respectively, at 4 °C overnight. β-actin or GAPDH was used as a reference. The membranes were washed three times with TBST buffer for 3 times, incubate with horseradish peroxidase-conjugated secondary antibodies at room temperature for 2 h. Detection was visualized using an enhanced chemiluminescence system. The protein signal was quantified using the ImageJ software.

### Quantitative reverse transcription PCR (qRT-PCR)

Total RNA was extracted from the BM, spleen, and blood using the conventional TRIzol method (Life Technologies, USA). The RNA precipitate was diluted 100 times with deionized water and then quantitatively analyzed using a spectrophotometer. Reverse transcription was performed using PrimeScript^TM^ RT Reagent Kit (Takara Bio Inc., Japan) with gDNA Eraser, in accordance with the manufacturer’s instructions for two-step reverse transcription polymerase chain reaction: first synthesis cDNA, and then PCR amplification was performed. Reverse transcriptase was omitted from the negative control to determine whether genomic DNA remained in the cDNA. PCR reaction system: SYBR ^®^ Premix Ex Taq (2 ×) 10 ul, cDNA 2 ul, 0.8 ul either upstream or downstream primers, ROX Reference Dye II (50 ×) 0.4 ul, steam sterilization double water 6 ul. Cycle parameters for: 95 °C pre-degeneration 30 s, into the PCR cycle at 95 °C degeneration 5 s, 60 °C annealing/extension in 34 s, for 40 cycles. The normalized expression values for each transcript were calculated as the quantity of target gene mRNA relative to the quantity of β-actin mRNA using the 2^−ΔΔCt^ method. All reactions were performed independently at least three times. The primer sequences used are listed in the Supplemental Tabel 1.

### Flow cytometry

The cell suspensions obtained from blood, BM, or human and mice spleens were submitted to flow cytometry analyses. Blood cells from human were incubated with APC-anti-CD19, PE-Cy7-anti-CD80, APC-H7-anti-CD86. Mouse BM cells were incubated with APC-anti-B220, PE-anti-CD138, and PE-Cy7-anti-IgM antibodies. Spleen cells from the mice were stained with combinations	of	MABs	specific	for APC-anti-B220, PE-anti-CD138, PE-Cy7-anti-IgM, and PE-anti-IgD. Spleen macrophages from mice were stained with APC-anti-F4/80, FITC-anti-CD11b, PE/Cy7-anti-CD206, and PE-anti-CD86. Blood cells from the mice were incubated with APC-anti-B220, PE-Cy7-anti-IgG, and PE-anti-CD138. All the antibodies were obtained from BD Pharmingen (San Jose, CA, USA). The samples were acquired on a BD Canton-II flow cytometer and analyzed using FlowJo software. Dead cell exclusion was performed using scatter profiles and 7-AAD staining during all the flow cytometric analyses.

### Immunofluorescence staining

Spleen sections were fixed in 4% paraformaldehyde for 30 min and then placed in a cassette filled with EDTA antigen buffer (pH 8.0) in a microwave oven for antigen repair. After shaking dry, the sections were placed in 3% hydrogen peroxide solution for closure and were then incubated with monoclonal rabbit anti-CD86 (Servicebio Technology Co., Ltd., Wuhan, China), monoclonal rabbit anti-CD206 (Servicebio Technology Co., Ltd., Wuhan, China), and monoclonal mouse anti-BAFF (Abcam plc., UK), followed by incubation with goat anti-rabbit/mouse corresponding species immunoglobulin antibodies. DAPI (Servicebio Technology Co., Ltd., Wuhan, China) was used to label nuclear DNA. Data of co-localization were analyzed by plugin ‘Coloc2’ of ImageJ software, and the Pearson’s R values selected above threshold.

### Statistical analysis

To examine the significance of the mean differences between groups, we used two-tailed unpaired or paired Student’s *t*-tests for normally distributed data and the nonparametric Mann–Whitney U test for non-normally distributed data. The relationship between the two features of interest was determined using the Spearman’s rank correlation test. All statistical tests were performed using GraphPad Prism V8.0. Unless otherwise specified. All *p*-values presented were two-sided, and statistical significance was set at *p* < 0.05.

### Study approval

For animal studies, experimental procedures were approved by the Experimental Animal Ethics Committee of Nanjing Medical University for the use of live animals for teaching and research. For human studies, the collection of peripheral blood was approved by the ethics committee of Nanjing First Hospital. Written informed consent was obtained from all patients and healthy donors.

## Author contributions

SL and XMM developed the study concept and directed experimental design with help from GQL, QWG, and JW. SL, GQL, QWG, JW, XC,YZ, JJM, QS and WSG performed experiments. SL, GQL, QWG, JW and XMM analyzed the data. SL and XMM wrote the manuscript. All authors read, edited, and approved the manuscript.

## Acknowledgements

This work was supported by National Natural Science Foundation of China (81570710). We thank the following core facilities of the Central Laboratory, Nanjing First Hospital, Nanjing Medical University, Nanjing, China for their assistance: Flow Cytometry, Cell Culture, Immunohistochemical, and Immunofluorescence.

---

**Supplemental Tabel 1.**
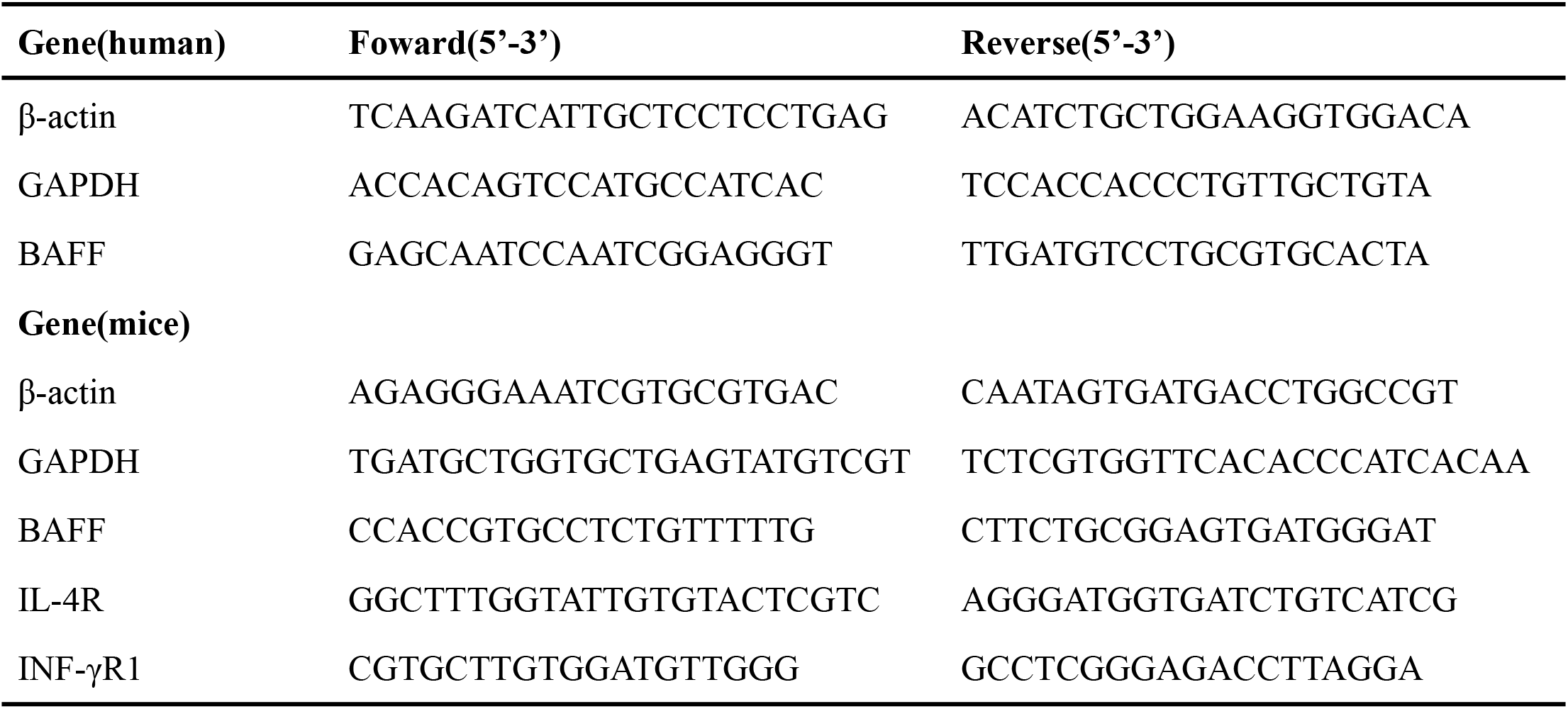
The primer sequences for RT-PCR in human and mice.

**Supplemental Figure 1.**
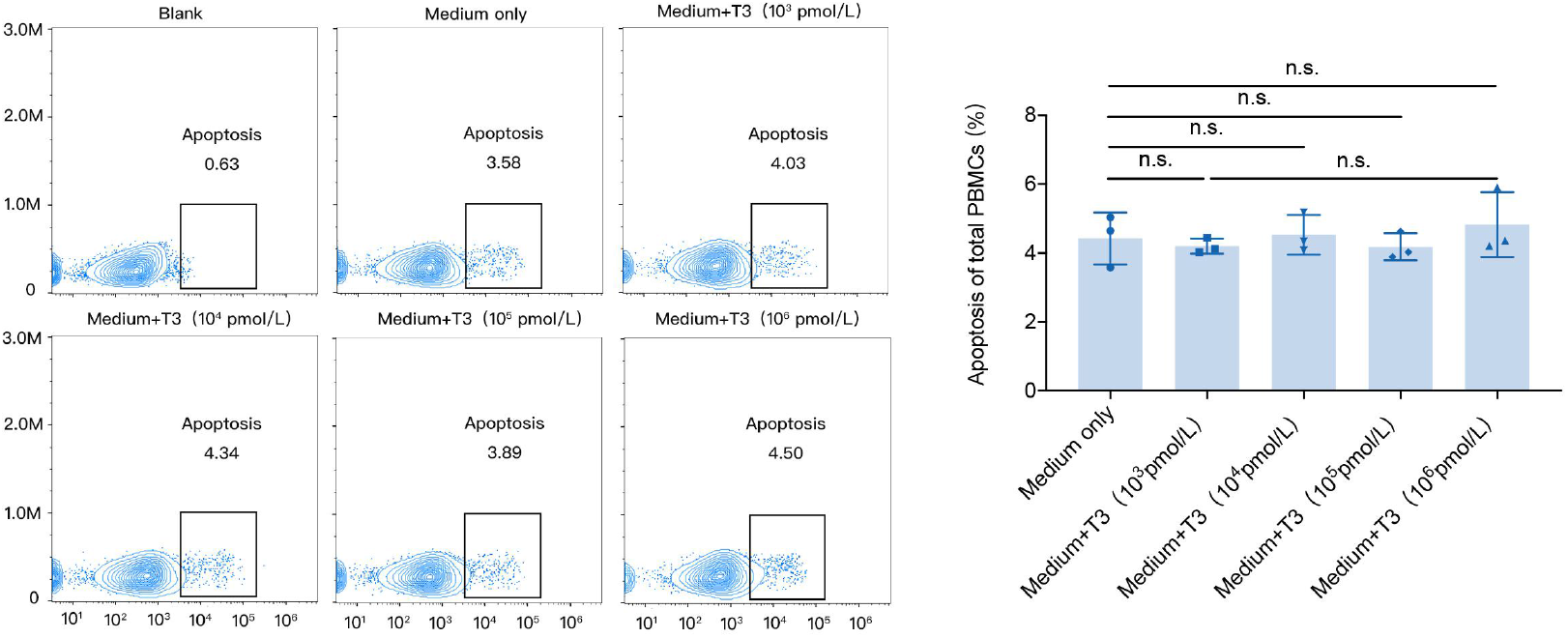
The apoptosis of PBMCs with various concentrations of T3 stimulation. The PBMCS were isolated from the normal mice and treated with various concentrations of T3 for 72h. The apoptosis of PBMCs was determined by staining with AnnexinV-PE. Data are presented mean ± S.D. Statistical significance is assessed by two-tailed unpaired *t*-test. (n = 3 independent biological experiments).

**Supplemental Figure 2.**
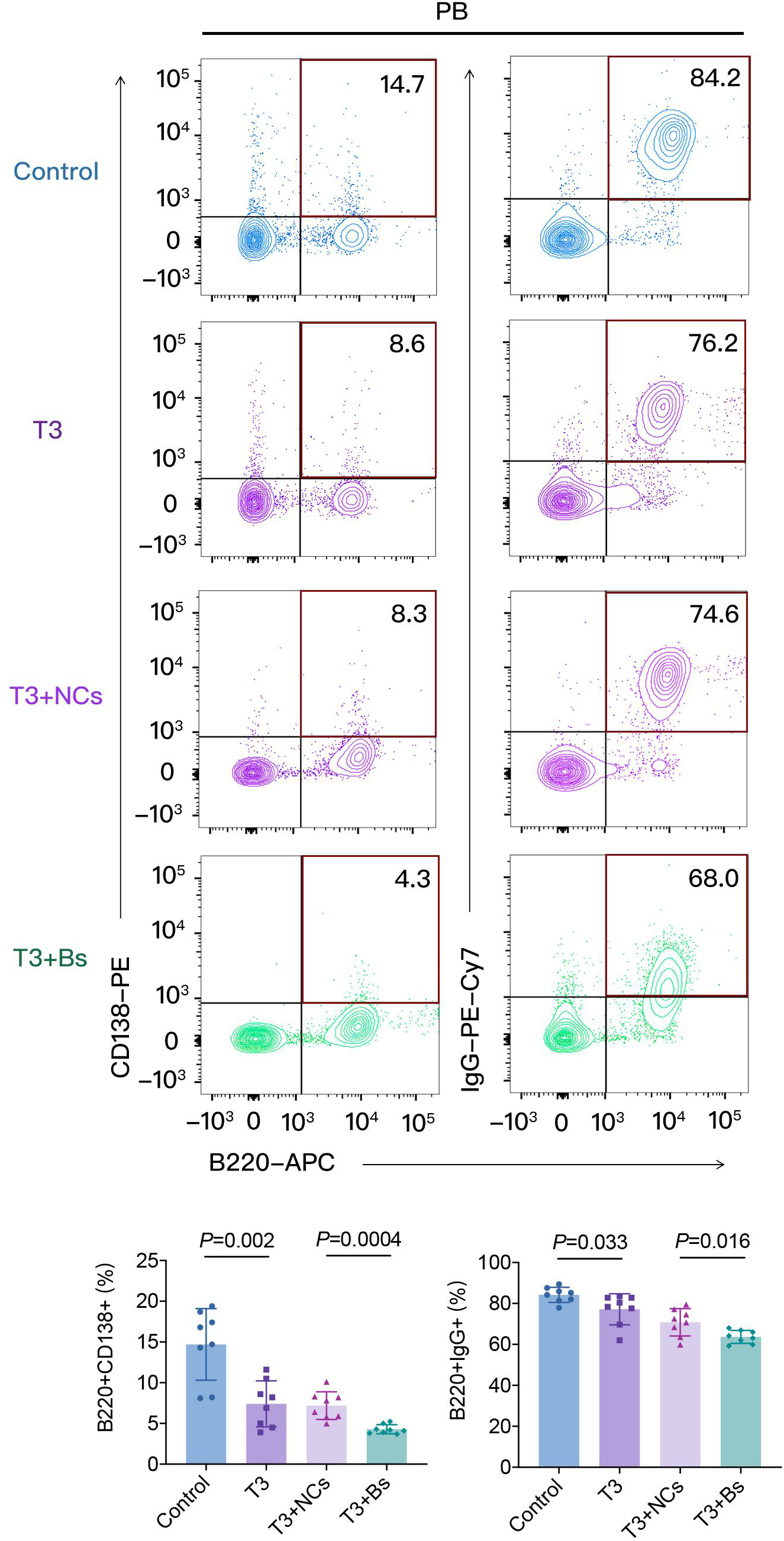
B cells differentiation in PBMCs of the mice after T3 treatment. Representative expressions of CD138 and IgG on B220+ B cells in PBMC of the control, T3, T3+NCs and T3+Bs groups. The average fraction of B220+ B cells positive for CD138 and IgG are shown at the bottom panel Data are presented as mean ± S.D. (n = 8 independent biological experiments). Statistical significance is assessed by two-sided independent *t* test.

