## Supplementary figures and images for "Thyroid hormone induces B cells abnormal differentiation via overexpression of B cell-activating factor"

### Supplemental Figure 1

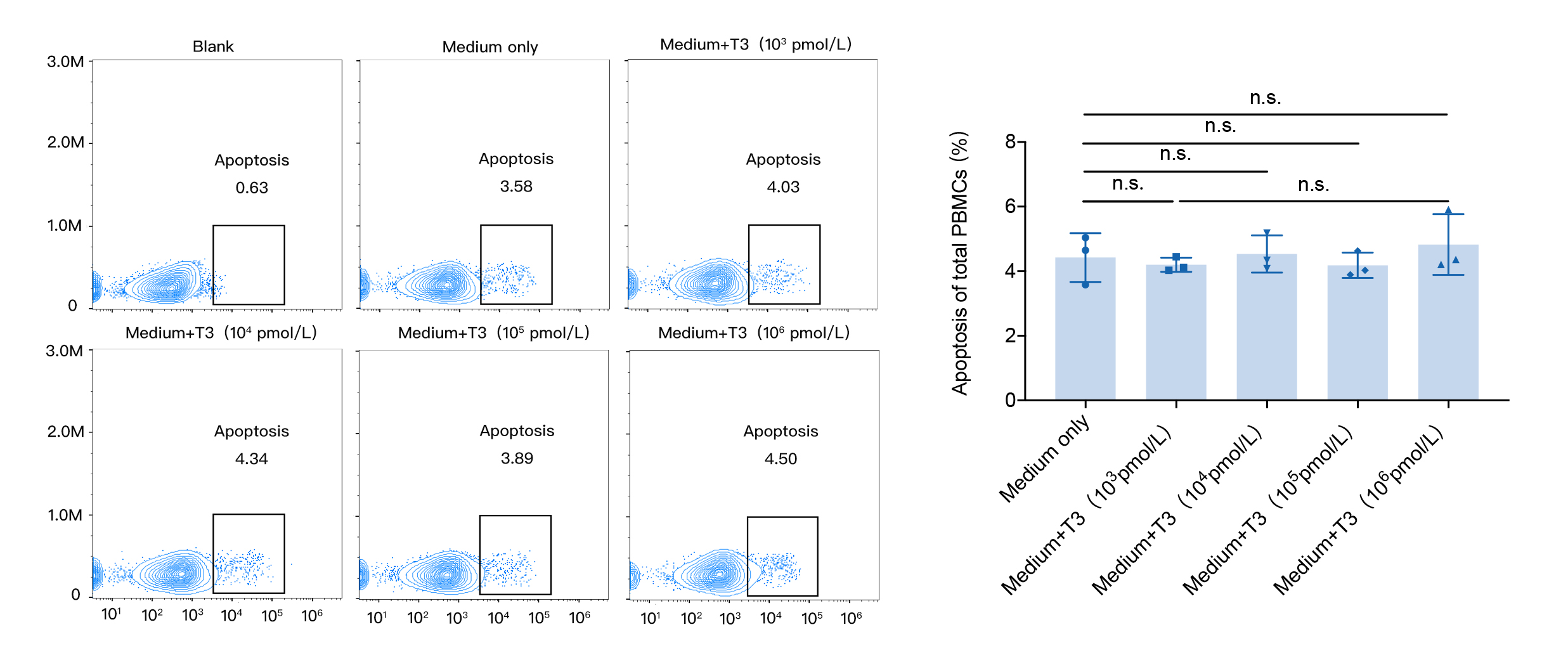
